# *EMS* hypothesis of Macroevolution

**DOI:** 10.1101/2025.09.24.678403

**Authors:** Xin-guo Yang

## Abstract

Starting from the unified topological geometric analysis framework of four ecosystem organization rules, including the Kleber’s 3/4 law, population -3/2 self-thinning law, ecological niche theory, and community neutrality principle, Ecosystem Metabolic Self-organization (EMS) is defined as a process in which individual or characteristic individual in group spontaneously construct their biological space in a three-dimensional resource space, within the redefined framework of ecosystem metabolic network. Under the conditions of metabolic steady (or metabolic conservation), resource diffusion dimension (D) and metabolic network dimension (D+1), which together define the biological space by at most three space parameters (x, y, z) and one direct parameter (d), unified the topological geometry principle and metabolic scaling function of the four rules, B∝M^D/D+1^.

The results indicated that EMS follows a Fibonacci sequence (*f*(n), n) evolution law at kingdom level, from neutral system (prokaryote 1, 1), niche system (unicellular eukaryote 1, 2), modular similarity system (plant 2, 3), to internal metabolic system (animal 3, 4), as indicated by (D, D+1). Further derivation revealed the positions of superbody system (fungi 5, 5) and primitive soup system (0, 0) in this evolutionary sequence. Thus, a conceptual model of macroevolution with the connotation of metabolism co-evolution between life and ecosystem was proposed.

This article took the four principles of ecosystem organization as theory prototypes and the analysis paradigm of biological space developed by metabolic theory of ecology (MTE) as referential methodology. It preliminarily constructed the logical, conceptual, and methodological system of EMS hypothesis, and provided a deducible Fibonacci roadmap for macroevolution. In any case, macroevolution should be simple and beautiful, just as the model of nature.

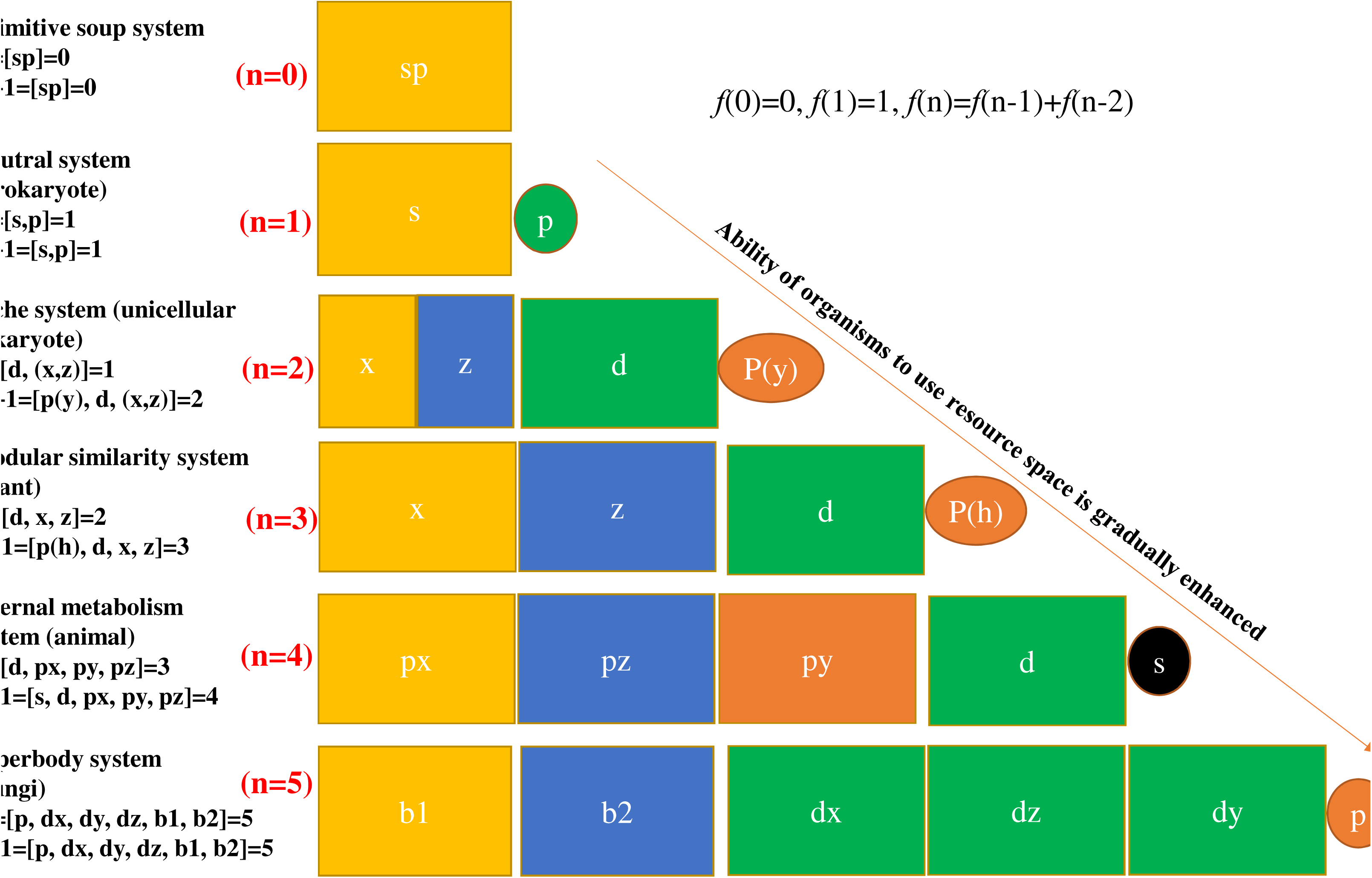

---

Kleiber’s 3/4 law (Kleiber, 1932), population -3/2 self-thinning law (Yoda et al., 1963), niche theory (Tilman, 1986), and community neutrality principle (Hubbell, 2001) potentially define ecosystem organizational rules at different levels from individual, population to community, involving resource utilization or metabolic activities. The four representative empirical models in fact constitute the bases of ecology. If there is a theoretical framework that can unify the four models, it will greatly enhance our understanding of the organization mechanisms of ecosystem.

This work was represented by the metabolic theory of ecology (MTE), which attempted to Unified Kleiber’s 3/4 law and -3/2 population self-thinning law (West et al., 1999a; Enquist et al., 1998), and applied to the general definition of ecosystem processes (Brown et al., 2004), based on the metabolic scaling theory (WBE model, West et al., 1997) under the geometric analysis paradigm of “biological space” (West et al., 1999b; Banavar et al., 1999). However, the uniqueness of the 3/4 rule across species had been widely questioned theoretically and empirically (Makarieva et al., 2008; DeLong et al., 2010; Hatton et al., 2019). By now, there have been no systematic attempts to unify the four empirical models, except for a recent study integrating individual and community metabolic scaling (Fant and Ghedini, 2024).

In order to build such a unified theoretical framework, **the choice of technical route should be considered firstly.** For example, biological geometry (Glazier, 2018) and metabolic topology (Ravasz et al., 2002) technical route have been tested to be suit. Based on the paradigm of biological space developed by MTE, however, it is necessary to extend these researches from life level to ecosystem level, that is, the ecosystem metabolic network research focusing on the resource utilization processes from environment to organisms.

First of all, from the perspective of the whole chain utilization of resources, ecosystem metabolism can be divided into two processes: internal metabolism and external metabolism. Internal metabolism is the second diffusion process after resources enter the body, while external metabolism is a process of resource capture and diffusion from the environment to body. Therefore, ecosystem metabolic network can be defined as a complete resource diffusion chain composed of the first diffusion process (external metabolism) and the secondary diffusion process (internal metabolism). Thus, it is logically feasible to extend from biological metabolic network to ecosystem metabolic network. Secondly, metabolic network topology research is a general analysis paradigm that focuses on the efficient use of resources and reveals how the topology optimization of network promotes the maximization of metabolic efficiency (Shellman et al., 2014; Morrison and Badyaev, 2016). For ecosystem metabolism network, it is to transform the bio-geometric behavior of resource utilization (Glazier, 2018) into the topological connection between biology and environment (the space of resource distribution). Therefore, the two could be unified in topology analysis.

Secondly, **the biological applicability of this unified theoretical framework needs to be limited.** Based on the research progresses in related fields, an empirical cognitive result can be drawn: the biological applicability of the four empirical models is significantly different. For example, Kleiber’s 3/4 law is very common in animals; Yoda -3/2 self-thinning rule is strictly applicable to modular organisms, especially photosynthetic multicellular plants. Although the classical niche theory (Tilman, 1986) and the community neutrality principle (Hubbell, 2001) are more proposed from large biology researches, they begin to get more and more applications in microorganisms with the progress of technology (Fierer and Lennon, 2011).

Studies have shown that the ecological characteristics of bacteria in high-order taxa are highly similar, and the local bacterial communities tend to show the characteristics of phylogenetic aggregation (Philippot et al., 2010), which is easier to meet the principle of individual ecological equivalence, so bacterial community is more suitable for the neutral principle. Compared with bacteria, there are fewer studies on the community construction of unicellular eukaryotic. But according to the principle of community construction proposed by weiher & Keddy (1995), there are strong interactions among species in the community with divergent phylogenetic relationships. The divergence of the phylogeny of the local community of unicellular eukaryotes is often more extreme (Vaulot et al., 2008; Burki, 2014), which is therefore more applicable to the niche principle. In fact, microbial community has been regarded as a potential ideal model to test the niche-neutral principles continuum.

If the specific historical background of niche and neutral principles research is ignored, an assumption can be drawn that the matching of ecosystem organization rules with specific life evolution types at different organizational levels (or organizational modes) may be the result of natural selection under macroevolution. The matching of neutral principle and prokaryote, as well as the combination of niche principle and unicellular eukaryote, conform to the general principle of macroevolution from simple to complex. **Thus, this general assumption also defines the macroevolution background of the application of the unified theoretical framework.**

Therefore, this theoretical framework needs to focus on the generalization of the rules of ecosystem organization under macroevolution, distinguishing from the current application of generalized niche theory and neutral principle, and strictly based on the biological or mathematical conditions of the four organization rules (i.e., metabolic steady defined in the article).

Accordingly, by integrating the ecosystem organization level, organization principle and applicable biological scope, this paper unifies the triple naming system for the four ecosystem organization modes. On this basis, this paper constructs an analytical framework of **EMS**, to unify the topological principles and functional expressions of the four modes, and its application in macroevolution. The basic concepts related to **EMS** can be seen in **concepts and methods I**.

## 1 “spatial embedding” conceptual model of EMS: united construction logic of biological space

To unify the construction logic of biological space for the four ecosystem organization modes, this paper proposes a spatial embedding conceptual model based on the construction of an ecosystem metabolic network in a three-dimensional resource space (Figure 1). Growth (or movement) serves as the fundamental mode through which organisms embed themselves into the three-dimensional resource space. Under the condition of metabolic steady, the four models are geometrically abstracted as the optimal embedding processes of a single point, solid a (homogeneous individual group), and a solid (heterogeneous individual group) into the three-dimensional resource space, thereby converting the problem into a topological geometry issue.

**Fig. 1.**
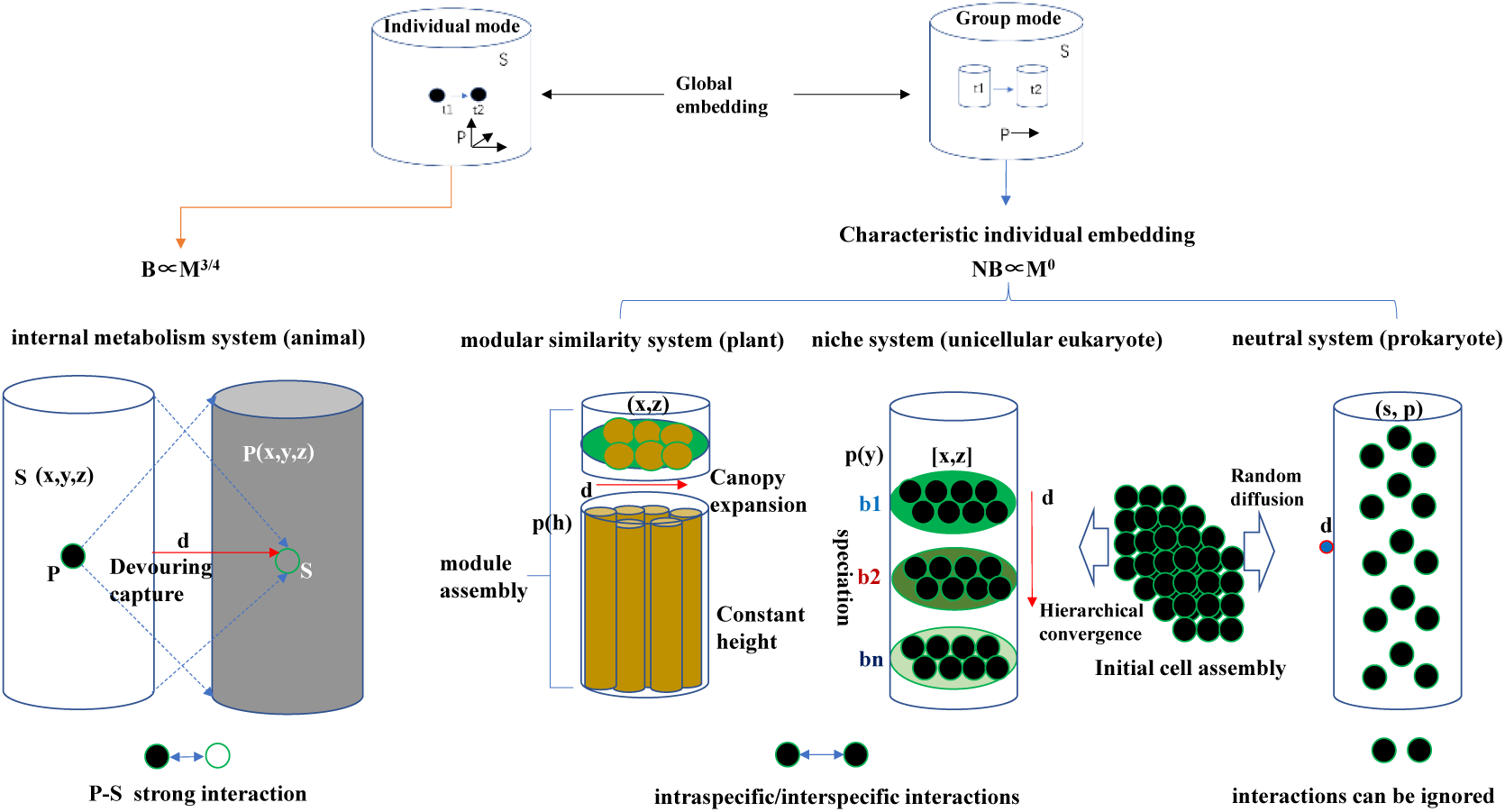
“spatial embedding” conceptual model and topological geometry principle of biological space construction. **S**, embedded space (environment); **P**, embedding unit (biology); **t1-t2**, Growth or movement of embedding units; **b1-b2-bn**, niche differentiation under the niche mode. **S (x, y, z)**, resource space 3D coordinate reference system; **P (x, y, z)**, three-dimensional coordinate reference system in animal body space; **d**, direction parameter (vertex) of space embedding (defining resource capture process, or position of biology individual at a time *t* after the embedding); **p**, biological individual vertex (embedding starting point); **p(h),** represents the starting state of the individual occupying the y-axis space with a fixed height h (population self-thinning mode); **p(y)**, represents the spatial distribution of individuals along the y-axis (niche differentiation axis) (niche pattern); **s**, resource space vertex (the whole resource space as a point)**. The same below**. Based on the biogeometric principles of the four ecosystem organizational modes: (1) For the individual-internal metabolism mode (animal), spatial embedding is defined as the overall inversion of the relative positional relationship between the resource space and the organism, which is caused by animal phagocytosis; meanwhile, the overall embedding relationship between a point and the three-dimensional space remains unchanged. (2) The population-modular self-thinning mode (plant) is defined by the self-thinning process driven by the geometric similarity of modular organisms, and spatial embedding is divided into the nesting of two modular processes. (3) The community-niche mode (unicellular eukaryote) is defined by the process of full differentiation and realization of potential niches in the resource space, and spatial embedding consists of the species differentiation process and the secondary aggregation process of conspecific individuals. (4) For the community-neutral mode (prokaryote), neutral mode (prokaryote), spatial embedding is defined by the random diffusion of individuals under conditions where inter-individual interactions are negligible.

Spatial embedding is divided into two modes: global embedding and characteristic individual embedding.

**Global embedding** describes the macro-topological relationship between a group (as the embedding unit) and the resource space. In the three-dimensional resource space, the organizational mode of a single individual is geometrically defined as a point topological unit—where the relative positional relationship between the organism and its environment is determined by three coordinate parameters. Populations and communities, by contrast, are defined as volume topological units— their relative positional relationships can be characterized by a single directional parameter (e.g., internal vs. external relations). Global embedding reflects the topological characteristics of the total mass distribution (NM) in the group’s organizational mode.

**Characteristic individual embedding** (i.e., spatial embedding shaped by characteristic individuals within a group) is an effective process for constructing biological space. This process fully accounts for differences in organisms’ metabolic behaviors and the intensity of interactions between individuals (e.g., intraspecific and interspecific competition)—factors that determine the topological structure of the ecosystem metabolic network and the realization of metabolic efficiency. The embedding of characteristic individuals reflects the topological characteristics of both the mass distribution (M) and metabolic efficiency (B) of individuals within the group.

## 2. Topological geometry model of EMS: unification of geometric principles by simplex topology

The major results of model are summarized as shown in Fig 1 and Fig 2. The concept and method description of topological geometry model in the **concept and method II**, and more analysis detail of different organizational modes can be seen in the **concept and method III**.

**Fig. 2.**
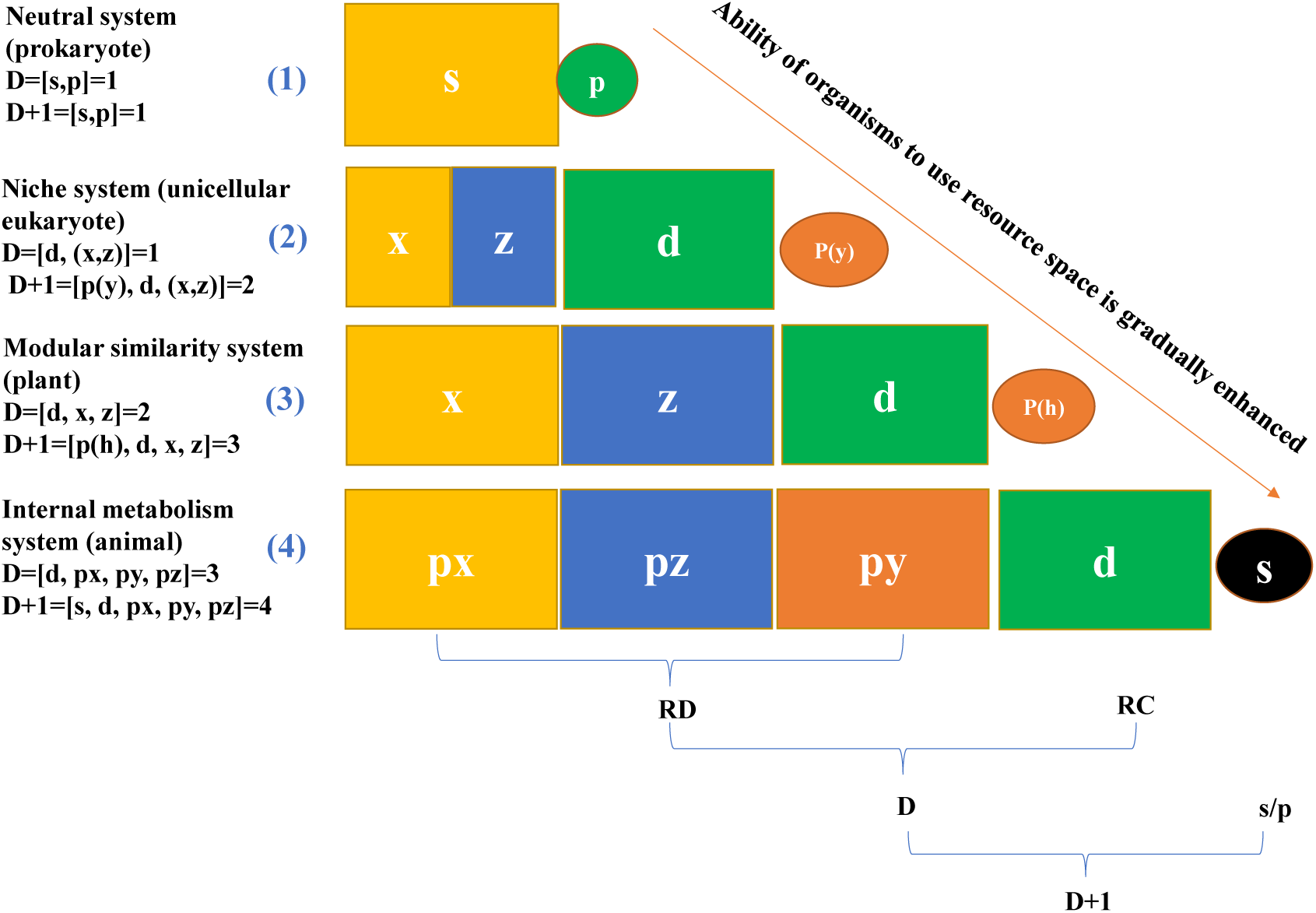
Origin and composition of the vertices of the D-simplex (resource diffusion dimension) and the D+1-simplex (metabolic network dimension). Under the unified scale constraint of topological space analysis based on the three-dimensional coordinate system of resource space, the construction of biological space (defined by resource diffusion dimension D and metabolic network dimension D+1) for the four organizational modes can be unified under a simplex geometric analysis framework. This framework consists of topological vertices involved in two spaces within the first resource process of the ecosystem metabolic network: the resource capture (or possession) behavior space (RC) and the resource diffusion (or utilization) behavior space (RD). These vertices include three-dimensional coordinate vertices (x, y, z), the embedded direction parameter vertex d, and the biological node p (origin). In the internal metabolic model (animal), the positional relationship between S and P is reversed.

In general, from the neutral system (prokaryotes) to the internal metabolic system (animals), the ability of organisms to utilize the resource space is gradually enhanced, and the topology of the ecosystem metabolic network becomes increasingly complex (Fig.2).

1. Internal metabolism system (animal). Resource diffusion is defined by three coordinate system parameters (py, px, pz) of the organism’s internal body space, while resource capture is defined by a d-s topological connection—this connection represents phagocytosis, a process that transfers resource points (s) into the organism’s body. The dimension of resource diffusion (D) is 3, as defined by the set [d, px, py, pz] (where d serves as the origin); the dimension of the metabolic network (D+1) is 4, as defined by the set [s, d, px, py, pz] (where s serves as the origin). Within this topological analysis framework, the fourth biological dimension of animals is clearly defined by the behavioral space of resource capture.
2. Modular similarity system (plant). The difference from the internal metabolic mode lies in that, due to the domain masking effect among individuals, the y-axis loses its indicative significance for resource diffusion of characteristic individuals and is only defined by the surface topological structure composed of the x-axis and z-axis. Under the constant constraints of h and metabolic conservation, resource capture (d-p(h)) is perpendicular to the y-axis. This is fully consistent with the biological geometric principle of population self-thinning.
3. Niche system (unicellular eukaryote). The y-axis, functioning as an ecological niche differentiation axis, has regained its spatial indicative significance but has been transformed into an indicator axis for resource capture activities (d-p(y)). Resource diffusion is indicated by the parameter space (x, z), which is defined by the combination of the x-axis and z-axis. Compared with the modular self-thinning mode, the aggregation space of conspecific individuals (depicted by the x-axis and z-axis) is constrained by the non-fixed random diffusion behavior of unicellular individuals, and their specific positions within this space cannot be determined. Topologically, this space is geometrically transformed into a vertex or a coordinate axis. This model is strictly restricted to conditions where potential ecological niches are fully differentiated within a limited space.
4. Neutral system (prokaryote). In the mode of completely random individual diffusion, resource diffusion and resource capture—defined by the relative positional relationship between P and S— hold no substantial topological significance. The three-dimensional coordinate reference meaning of the resource space is entirely lost, and the space is geometrically defined as a single vertex s. The construction of biological space is solely defined by passive resource diffusion driven by the environment (s-p). Under conditions of completely random diffusion, the organizational patterns of bacterial populations or communities have no impact on the outcomes.

In conclusion, the topological geometric analysis paradigms for biological space construction under the four modes are completely unified: all of them are topological space definitions for two types of biological geometric behaviors—resource capture and resource diffusion—centered around three spatial coordinate parameters and one spatial embedding direction parameter. The differences among the four modes can be accounted for by the changes in the parameter sources and topological properties that define resource capture and resource diffusion respectively.

## 3 Metabolic scaling model of EMS: unification of functional form

On the premise of metabolic conservation (NB∝M⁰), with M serving as the intermediary for the geometric transformation from the topological model to the scaling model, the unified metabolic scaling model is derived by constructing the topological dimensional relationship between M and NM (table 1). The metabolic scaling model of EMS is as follows,

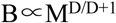

**Table 1.** Conversion and derivation from topological geometry model to metabolic scaling model.

| Conversion/<br>derivation | $NB \propto M^0$ | | |
| --- | --- | --- | --- |
|  | Modular similarity system | Niche system | Neutral system |
| 1). $D_M = D+1$ | 3 | 2 | 1 |
| 2). $D_{NM}$ | 1 | 1 | 0 |
| 3). $NM \propto M^a$ | $NM \propto M^{1/3}$ | $NM \propto M^{1/2}$ | $NM \propto M^0$ |
| 4). $B \propto M^b$ | Due $NB \propto M^0$, $NM \propto M^{1/3}$<br>$\Rightarrow B \propto M^{2/3}$ | Due $NB \propto M^0$, $NM \propto M^{1/2}$<br>$\Rightarrow B \propto M^{1/2}$ | Due $NB \propto M^0$, $NM \propto M^0$<br>$\Rightarrow B \propto M^{1/1}$ |
| 5). $B \propto M^{D/D+1}$ | Due $b=2/3$, $D=2$, $D+1=3$<br>$\Rightarrow B \propto M^{D/D+1}$ | Due $b=1/2$, $D=1$, $D+1=2$<br>$\Rightarrow B \propto M^{D/D+1}$ | Due $b=1/1$, $D=1$, $D+1=1$<br>$\Rightarrow B \propto M^{D/D+1}$ |

Note: $D_M$ and $D_{NM}$ represent the dimensions of their respective mass distributions in the three-dimensional resource space. $D_{NM}$ is defined by the global embedding mode, $D_M$ is defined by the characteristic individual embedding mode.

B, characteristic individual metabolic efficiency; M, characteristic individual mass; D, resource diffusion dimension; D+1, metabolic network dimension. The transformation logic and derivation process can be seen in the **concepts and methods IV.**

It can be observed that the macro-topological structural characteristics of the ecosystem metabolic network define the power function relationship between the metabolic rate and the mass of characteristic individuals. This relationship reflects the macro-constraints imposed by the group living mode on individual metabolism under metabolic conservation, and indicates the ecosystem attributes and compositional sources of the metabolic index *b* (where B∝M*^b^*). This also applies to animal individuals when resource capture in the resource space is taken into account.

## 4 Deduces

### 4.1 Fibonacci evolution model of EMS

The macroevolution of EMS is regarded as a continuous change process consisting of the steady state (point) and transition state (interval) of metabolism. The Fibonacci sequence discrete function defined by the four organization modes (under the steady state point) depicts the profile and trend of EMS macroevolution at kingdom level (Fig. 3). The definition of transition state indicates the potential existence of non-integer power-law for EMS at species or cross species scale.

**Fig. 3.**
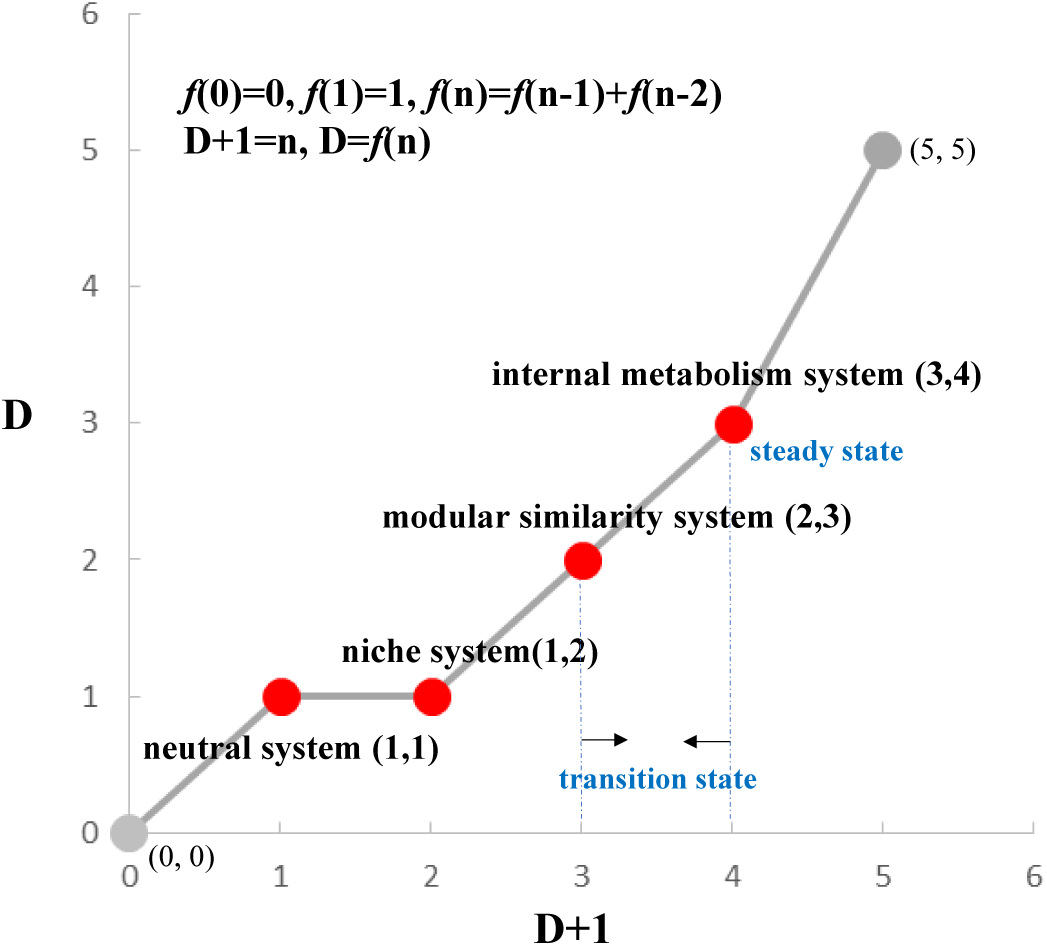
Fibonacci evolution model of EMS

This indicates a potential constraint mechanism for the evolution of EMS: Golden Section. The corresponding deduce is that the construction of biological space under the four organizational modes can essentially be regarded as the process of optimizing the use of space based on the golden section. The Fibonacci sequence evolution model of **EMS** can be preliminarily constructed as following,

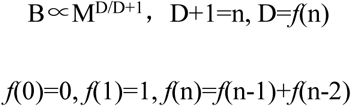

In this model, superbody system (i.e., mycelium fungal; an ecosystem metabolic network composed of mycelium fungal and three-dimensional resource space) and primitive soup system (the primitive ecosystem metabolic organization type before the emergence of cell life forms) are the other two potential macroevolution sequence objects. Applying the unified topological geometry principle of EMS, it is further deduced and established the status of superbody system (*f*(5)=5) and primitive soup system (*f*(0)=0) in the Fibonacci sequence (detail of analysis can be seen in **concept and method II** and **Fig.S1 in Supplementary materials**)

### 4.2 Applicable biological classification scope of EMS

Based on the derivation results of EMS topological geometry model and Fibonacci model at kingdom level, the applicable biological taxonomic scope of EMS can be further refined at phylum level (or equivalent taxonomic group), taking five kingdom classification as a reference.

The results are as follows: primitive soup system (0, 0), neutral system (prokaryote/***bacillus*** 1, 1), niche system (unicellular eukaryote/***diatom*** 1, 2), modular similarity system (plant/***photosynthetic multicellular plant*** 2, 3), internal metabolism system (animal/***internal circulation animal*** 3, 4), and superbody system (fungi/***mycelium fungi*** 5, 5). It can refer to the relevant summary and discussion section, as well as Table.S1 in Supplementary Materials.

### 4.3 conceptual model of macroevolution: synergy between ecosystem evolution and life evolution

From the primitive soup system to the superbody system, organization degree (topological complexity) of ecosystem metabolic network is gradually improved, consistent with the trend of complexity of biological metabolism. By EMS as an indicator of macroevolution, this paper proposes a macroevolution conceptual model: co-evolution process between life evolution forms and ecosystem metabolic self-organization modes with the goal of maximizing metabolic efficiency (Fig. 4).

**Fig. 4.**
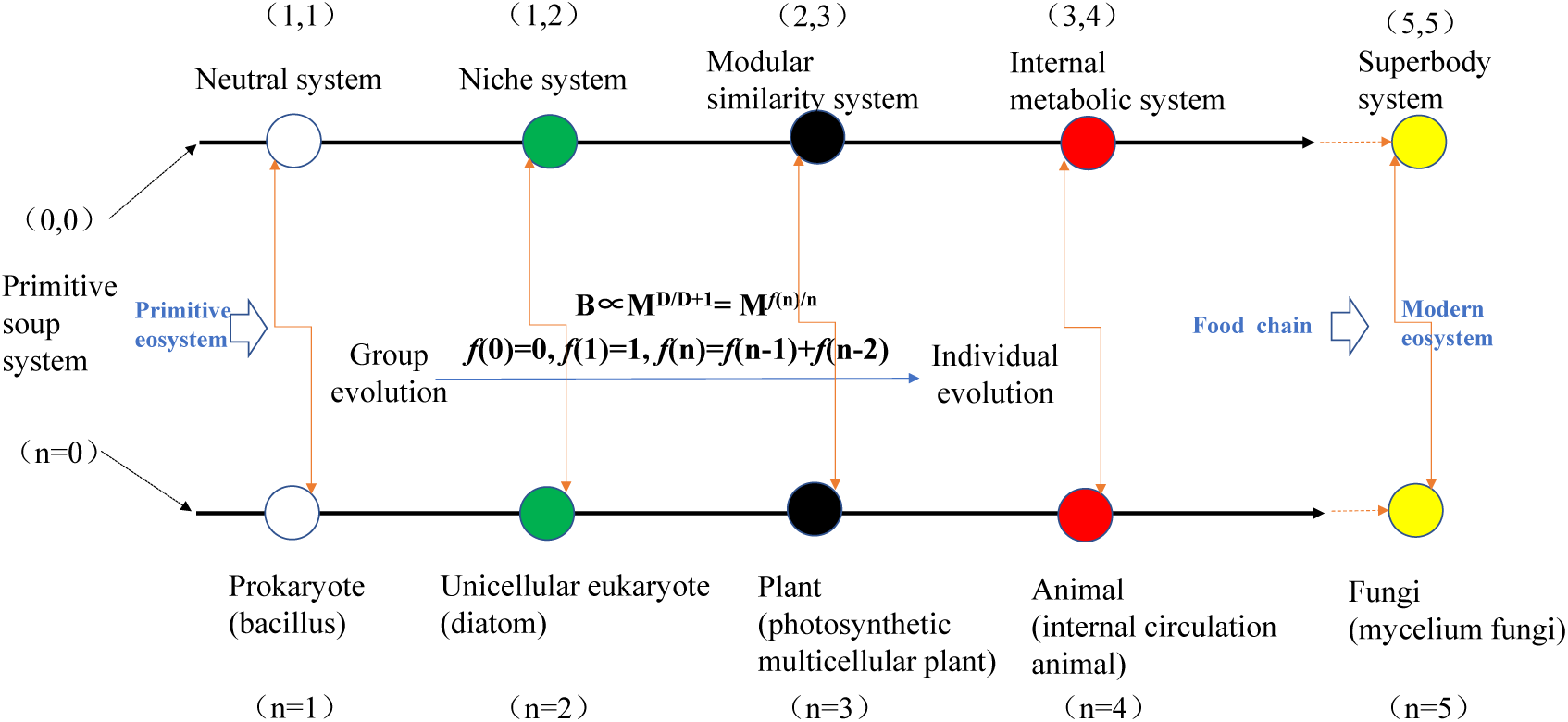
Conceptual model of macroevolution (cellular life). The order at kingdom level has nothing to do with the occurrence time of different life types, but only reflects a macro evolutionary logic based on structure and function, similar to the earliest five kingdom classification.

The macroevolution logic of the co-evolution model is that, the primitive soup system provides the primitive ecosystem environment for the breeding of cell life, the neutral system is the embryonic form of ecosystem evolution, and the niche system has spawned the most important and basic process of life diversity differentiation in the history of life evolution. On this basis, plants (modular similarity system - full use of light energy), animals (internal metabolic system - flexible predator-prey regulation), and mycelium fungi (superbody system - efficient waste decomposition) have taken on the important roles of producers, consumers and decomposers in the modern ecosystem with their own specialized ecosystem metabolic network, thus completing the evolution from the primitive ecosystem to the modern ecosystem.

## 5 Summary and Discussion

### 5.1 EMS provides a holistic logic for macroevolution

Taking the four principles of ecosystem organization as prototypes and the biological spatial analysis paradigm developed by MTE as reference, this article has preliminarily constructed a logical, conceptual, and methodological system for EMS. The difference between EMS and MTE lies in the following aspects: first, the metabolic index b is not unique in EMS; second, the four metabolic indices constructed in EMS are based on the four ecosystem organizational rules, uniformly derived from the topological geometry principles of the ecosystem metabolic network, and conform to the discrete distribution law of the Fibonacci sequence. On the whole, EMS has broken through the MTE framework (West et al., 1997; Glazier, 2018).

And, the results derived from EMS are more in line with the overall evolutionary trend of the cross-species metabolic index *b* (Makarieva et al., 2008; DeLong et al., 2010; Hatton et al., 2019). The detailed discrepancies from empirical research findings can be attributed to three factors:

Scale differences: Disparities in kingdom or phylum levels ***versus*** cross-species levels based on individual species;

Principle differences: Topological geometric principles of ecosystem metabolic networks ***versus*** biological geometric principles at the organismal level;

Source differences: Model-derived metabolic topological behaviors of individual organisms or characteristic individuals within groups ***versus*** measured metabolic ratios of independently living individuals.

In any case, the metabolic index *b* in EMS functions as an indicator of macroevolution, rather than a definition of metabolic efficiency for a specific species or population. Fossil research evidence from Payne JL, et al. (2009) suggests that the first sudden increase in cell size over the 3.5-billion-year evolution of life occurred during the stage when eukaryotic cells emerged. By reversing the metabolic scaling of unicellular eukaryotes in the niche mode (M∝B²), it can be observed that the relative growth efficiency of unicellular eukaryotes is the highest among all four modes—that strongly corresponds to the aforementioned fossil evidence, rather than the empirical results (DeLong et al., 2010). The niche model also clearly demonstrates the important and fundamental role of the unicellular eukaryotic life stage in the cultivation and differentiation of biodiversity during macroevolution. This constitutes a significant difference between EMS and MTE in their application to macroevolution.

Similarly, the clustering (Philippot et al., 2010) or divergence (Vaulot et al., 2008; Burki, 2014) of phylogenetic characteristics in local microbial communities, as well as their adapted ecosystem organizational modes, are closely associated with their respective macroevolutionary backgrounds. Furthermore, the positions and logical roles of animals (internal metabolic system), plants (modular similarity system), and mycelial fungi (hyperbody system) in the macroevolution of ecosystems are clear, and the evolutionary clues regarding metabolic topological behaviors among these three groups are also well-defined. The position of the primitive soup system in the Fibonacci evolution model and its relationship with the neutral system (prokaryotes) further indicate that the primitive soup hypothesis is valid, and the five-kingdom classification system of cellular life remains reasonable.

EMS provides a holistic logic for macroevolution, especially considering the opposition and unity between complex self-organization (Kauffman, 2002) and natural selection: the objects of natural selection are concentrated at the ecosystem level, while complex self-organization is explained by life activities themselves, and then the opposition between the two is unified through the macro-synergy of ecosystem evolution and life evolution in terms of metabolism.

This holistic logic can be further elaborated from the evolutionary relationships among the first resource diffusion (external metabolic network), biological nodes, and the second resource diffusion (internal metabolic network), which together defining the ecosystem metabolic network (for more details, see **Table S1** in the Supplementary Materials).

### 5.2 EMS provides an analysis paradigm of biological space under macroevolution

An interesting phenomenon in the evolution of biological morphology is the predominance of bacilli (exhibiting one-dimensional growth) among prokaryotes, as same as diatoms (exhibiting two-dimensional growth) among unicellular eukaryotes, in contrast to the common spherical cells. However, this phenomenon can be uniformly explained by the topological geometric principle underlying biological space construction, for characteristic individuals in group under the constraint of metabolic conservation.

Compared with independently living individuals, group living can significantly influence individual growth and metabolic behaviors (Fant & Ghedini, 2024). In the early stages of life evolution, the critical mode of group metabolic activity that restricts phenotypic evolution may have been constrained by the scarcity of suitable living environments and resources—and the probability of this mode occurring should have been relatively high. Subsequently, this phenomenon has been preserved to the present day through the genetic fixation of morphology in certain species. Similarly, this result does not emphasize the generalization of topological configurations at the species level. Nevertheless, there is no doubt about the taxonomic status of bacilli and diatoms at the level of their respective kingdoms or phyla, as well as their important roles in ecosystem. The same holds true for photosynthetic multicellular plants, animals with internal circulation, and mycelial fungi. It is reasonable to conclude that the significance of EMS as an indicator of macroevolution is valid. In summary, the basis of natural selection lies in biodiversity, yet the outcome of natural selection—i.e., the specific type of life evolution that ultimately drives macroevolution—is unique.

In addition, by integrating the biological scale concepts (biological length *l* or biological time *t*), *l* or *t* of characteristic individuals in group is topologically equivalent to the spatial embedding direction defined by d-p. When extended to individual animals (*l*, or *t*∝M^1/4^), this refers to the one-dimensional movement of individual animals for resource capture, i.e., the fourth biological dimension (West et al., 1999b). This is consistent in connotation with the 1/4 power law of biological time (Burger et al., 2021): *life lies in motion*. Similarly, this biological length or time for mycelial fungi may correspond to a three-dimensional resource capture behavior (*life lies in occupying l*, or *t*∝M^3/5^), but it is dimensionless for prokaryote individuals in group under metabolic conservation (*life lies in freedom l*, or *t*∝M^0^).

In general, biological space can be understood as a metabolic topology structure characterized by the mass distribution (either through growth or movement) of organisms in the three-dimensional resource space, in order to the optimized utilization of resources. Within the framework of EMS, both the ecological origins of biological space—specifically, the interactions between organisms and their environment (including other organisms)—and its geometric origins—namely, the topological construction of ecosystem metabolic networks—are clearly defined. Moreover, these origins can be unified across different ecosystem organizational modes (as well as different life evolution types) under macroevolution.

### 5.3 EMS provides a deducible Fibonacci roadmap for macroevolution

The Fibonacci sequence can be traced back to the “rabbit sequence” proposed by Fibonacci in 1202, and it serves as an infinite approximation function for the golden ratio point. A prominent characteristic of this function lies in its “inherited” summation operation, with the first two numbers of the sequence being directly assigned values.

The logic behind its application to macroevolution is as follows: First, the evolution of life exhibits a distinct inheritability. The establishment of the macroscopic direction of life evolution is marked by the first and second endosymbiotic events (Martin et al., 2017), which represent a typical “summation operation” in metabolic evolution. Second, based on the unified principles of topological geometry, this article clarifies the initial evolutionary states of the primitive soup system (where *f*(0) = 0) and the neutral system (where *f*(1) = 1). This also validates the general hypothesis regarding the early life evolution.

If we compare macroevolution of cellular life to rabbit reproduction and measure it by the evolution of metabolic systems, the primitive soup system represents the developmental stage of cellular life, while the neutral system (prokaryotes) corresponds to the maturation stage of cellular life. The niche system (unicellular eukaryotes) is analogous to the first “self-reproduction” of cellular life (i.e., the first endosymbiosis), and the second endosymbiosis lays the evolutionary foundation for the modular similarity system (plants). After the secondary endosymbiosis, for photosynthetic organisms, the fixed growth mode, modular growth mode, and gregarious living pattern undoubtedly confer natural selection advantages—these traits facilitate maximizing metabolic efficiency centered on photosynthesis.

In fact, secondary endosymbiosis does promote the evolution of the hierarchical topology of the cellular metabolic network in unicellular algae (García et al., 2023), and the hierarchical topology just is similar to the macro-topology of the ecosystem metabolic network of the modular similarity system (Fig. 1). This is an interesting topological self-similarity, which is also reflected in other ecosystem organization and life types (**Table S1** in the supplementary materials).

So, is there a third or fourth endosymbiotic event that laid the early evolutionary foundation for the internal metabolic system (animals) or the superorganism system (mycelia fungal), respectively? Currently, there is no conclusive evidence. Further answers may be sought from the following aspects: the fuzzy boundary properties in the taxonomy of protophytes, protozoa, and protofungi; the discovery of plant-animal transitional life forms in the evolutionary stage; and the early evolutionary relationships between fungi, animals, and plants (Hugoson et al., 2022; Song et al., 2024). The evolutionary stage of Protista is a period when life forms and functions were highly diversified, which can be likened to a “**plasticine**” experiment of metabolic networks shaped by natural selection, from the perspective of the geometric nature of topology. Perhaps the answer lies within this process.

The EMS hypothesis provides a deducible Fibonacci roadmap for macroevolution, and this roadmap applies not only to the past. Similarly, assumptions can also be made about the evolution that go beyond cellular life. For instance, there are the virus and virus-host system (*f*(6)=8), as well as the nervous system and neuro-human body system (*f*(7)=13). Studies have revealed that the brain’s nervous system can operate across up to 11 topological dimensions (Reimann et al., 2017). It is speculated that within the context of the neuro-human body system, this dimension should be 13—adding two dimensions derived from the body surface? From the perspective of visualizing of biological space, it is further demonstrated that the underlying key elements are: the hexagonal inner shell of the phage capsid (which serves as the geometric evolutionary steady-state solution for the protection and release of nucleic acid), and the neural network (where seven network modules, or computing units, are cascaded to achieve complex perceptual tasks).

Compared with cellular life, the evolution types of life at this stage do not have independent basic metabolic capacity, and are more dependent on their ecosystem. The metabolic dimension D_B_ is larger than the mass dimension D_M_, which reflects the higher efficiency of ecosystem construction and the relative metabolic benefits of individuals (subsystems). Under the background of macroevolution, the co-evolution history between virus and bacteria is the longest, and it is a typical representative of early ecosystem regulation mechanism. The nervous system is the representative of wisdom, which embodies a higher level of ecosystem regulation mechanism. The metabolic activities of such systems are essentially carried out around “information”. Compared with “resources”, it is a higher level of ecosystem metabolic network evolution stage, and it is also the foreshadowing buried in the cellular life evolution stage: the transition of the complexity of the network structure inevitably requires the replacement of the system regulation mode. Therefore, a further speculation is that AI and AI-human society (Internet) system (*f*(8)=21), an evolution stage of swarm intelligence.

In any case, EMS hypothesis provides a global evolution framework for both cellular life and non-cellular life. Macroevolution should be simple and beautiful, just as the model of nature. Compared with the Red Queen model (Van Valen, 1973) and the court clown model (Benton, 2009), EMS endows macroevolution with an integrated logic of life and environment, and a deducible Fibonacci road map. The opposition between the complex self-organization (Kauffman, 2002) and natural selection can also be unified in the co-evolution framework of ecosystem and life. Therefore, the unification of the theoretical framework of macroevolution can be expected (Mitchell, 2009).

## Acknowledgment

Thanks to Professor *Wang Yong-xi* (Northwest University of agriculture and forestry science and Technology), Professor *Cheng Xu* (China Agricultural University) and Professor *Song Nai-ping* (Ningxia University) for their help in the research.

Concepts and Methods

## I Basic concepts of EMS

1. **Ecosystem metabolic network**: ecosystem metabolic network is a macroscopic topological connection network consisting of resource utilization activities between organism and its environment, composed of the first resource diffusion from environment to body (external metabolism) and the second resource diffusion in body (internal metabolism) activities. The first diffusion is composed of resource capture (or occupying) and resource diffusion (or utilization). With the aim of maximizing metabolic efficiency, organisms tend to establish universal connections (based on growth or movement) with three dimensions of resource space, achieving the goal of fully occupying and utilizing resource space.
2. **Environment (three-dimensional resource space):** Starting from the macroscopic topological connection between organisms and the environment, the environment is defined as a resource space with clear boundaries, defined by three spatial coordinate system dimensions (x, y, z). In topological geometry, it is defined as a structured spatial node that serves as a basic constraint for the macroscopic topological structure reference frame of ecosystem metabolic network.
3. **Ecosystem metabolic self-organization:** The process by which organisms (individuals or character individuals in group) spontaneously construct their biological space within a three-dimensional resource space. Within the framework of ecosystem metabolic network, biological space can be understood as a macroscopic topological structure driven by the optimized distribution (input) of biological mass in the three-dimensional resource space during the first resource diffusion. The macroscopic topological structure firstly is defined by the relative position relationship between organisms and resource space. Except for animal, the second resource diffusion is regarded as a black box, topologically equivalent to a biological node for the biological space.
4. **Ecosystem metabolic self-organization mode**: There are four types of ecosystem self-organization modes at kingdom level: individual-internal metabolic mode (animals), population-modular self-thinning mode (plants), community-niche mode (unicellular eukaryotes), and community-neutral mode (prokaryotes). The corresponding types of self-organizing ecosystem is named as internal metabolic system, modular similarity system, niche system, and neutral system, respectively.
5. **Metabolic steady**: An overall stable state or organizational condition that maximizes metabolic efficiency in the metabolic network of an ecosystem. Metabolic steady is a strict constraint for biological spatial construction under different ecosystem organizational patterns.

From perspective of generalization of the four ecosystem organization rules, following the principle of equivalent transfer of metabolic steady, the metabolic conservation (NB∝M^0^) defined by self-thinning populations is applicable to all group modes (population and community). Similarly, there exists a power function relationship between the metabolic efficiency and mass of characteristic individual in group (B∝M*^b^*), formally consistent with animal.

See separately, the condition corresponding to metabolic steady is, the occurrence of population self-thinning trajectory lines for the population-modular self-thinning mode (plants), total potential niches are fully occupied (the necessary condition of metabolic conservation) for the community-niche mode (unicellular eukaryotes), individuals are completely randomized in the resource space for the community-neutral mode (prokaryotes), and mature state of circulatory system for the individual internal metabolic mode (animals), respectively.

## II Concept and method of the topological geometry model of EMS

Convert the spatial embedding concept model into a topological geometry language, where the resource diffusion dimension (D) and metabolic network dimension (D+1) are used to define the overall macroscopic topological behavior of biological spatial construction. And it is defined that,

### Embedding starting point

It is defined by the embedded biological or ecological preconditions, that is, the original tenability conditions of the theoretical model corresponding to the four organization modes, and metabolic conservation is followed in group mode.

### Embeddedness drive

It is jointly defined by the relative position relationship between individuals and resource space, as well as the interaction between individuals (intraspecific or interspecific competition).

### Embedding trajectory

That is, the construction process of biological space, which is constrained and defined by the topological behavior of biological growth (or movement) in the three-dimensional resource space.

1. **Embedded space:** denoted as S, a Euclidean geometric space R³(resource space). In spatial embedding mode, it is the constraint parameter that determines the macroscopic topological structure characteristics of the ecosystem metabolic network. Under initial conditions, it is assumed to be a homogeneous space where available resources are uniformly distributed along three coordinate axes. During the process of spatial embedding, driven by biological behavior, the distribution attributes of homogeneous resources are altered.
2. **Embedded unit:** denoted as P (individual or characteristic individual in group). Compared to the three-dimensional resource space, in the global embedding mode, individuals are geometrically positioned as a point topology unit (with positional relationships defined by three coordinates), while the group is geometrically positioned as a body topology unit (with positional relationships defined by a directional parameter). In the metabolic network of ecosystems, P is both the starting point of spatial embedding and the endpoint of the first resource diffusion.
3. **Resource diffusion dimension:** denoted as D, it defines the topological dimension of the first resource diffusion activity in ecosystem metabolic network. It is defined by the parameter space consisting of the minimum number of parameters that describe the relative position relationship between P and S during the spatial embedding process. In the three-dimensional resource space, the minimum parameter for the relative positional relationship between P and S is defined by three coordinate points (x, y, z), and an embedding direction auxiliary descriptive parameter d is introduced. The connection between d and p is used as a directional description of the dynamic embedding (growth or movement) of individual. In the case where p is defined as the embedding starting point and serves as the topological origin, the basis for the existence of d as a vertex is the relative position descriptive parameter p_t_ of p at a certain time *t* during growth or movement. In the individual internal metabolism (animal) mode, the spatial positions of P and S are reversed.

Topologically, D is defined as a topological simplex consisting of coordinate parameters and directional parameters connected as vertices. D determines the fundamental topological properties of biological resource utilization activities proportional to the efficiency of biological metabolism. Taking animals as a reference, D can also be referred to as a metabolic dimension.

Resource capture (possession) and resource diffusion (utilization) constitute all two links of the first resource diffusion process in ecosystem metabolic network, and are also the two major sources of topological spatial activities that make up D. Resource capture (possession) is defined by biological growth (movement) activities, namely d-p topological connections. Resource diffusion (utilization) is defined by the static positional relationship between P and S, with three coordinate axes (x, y, z) serving as a universal reference frame in the three-dimensional resource space. The relative effectiveness of resource utilization for organisms, through resource diffusion activities indicated by different coordinate axes, determines the number and composition of parameters required to describe the static positional relationship between P and S under different organizational modes.

(4) **Metabolic network dimension:** denoted as D+1, which is the overall topological dimension of the ecosystem metabolic network (i.e. biological space dimension. Compared to D, it can be referred to as a structural dimension, while D is referred to as a functional dimension), defined by the minimum embedding spatial dimension containing P and S. The overall topology of the ecosystem metabolic network is the biological space defined by by the embedding trajectory of characteristic individuals, which includes both resource capture and resource diffusion.

D+1 is defined as a simplex constructed by P as the origin (embedding starting point) and all vertices in D. The construction of biological space can be concretized as the topological distribution of biological mass in three-dimensional resource space, which is proportional to the size of biological mass. Taking animals as a reference, the dimension of metabolic networks can also be referred to as a mass dimension.

### Additional explanation

In the spatial embedding mode defined by the characteristic individual P and the three-dimensional resource space S, the analysis scale of macroscopic topological structure of ecosystem metabolic network is uniformly defined by the constraints of the three coordinate dimensions of the resource space. Under this scale constraint, the resource utilization activities of characteristic individuals are dominated by intra- or inter-species competition in the first resource diffusion process, and the overall metabolic activity within organisms is regarded as a black box, topologically indicated by P as a vertex. In the individual internal metabolic pattern (animal), the reversal of the relationship between organisms and the environment is determined by the utilization of animal resources, and both primary and secondary resource diffusion activities are considered simultaneously. In summary, if the phagocytic behavior of animals is topologically equivalent to the displacement effect of resource space, and the internal circulation system is functionally equivalent to replacing the resource diffusion process in the first resource diffusion process, the construction of biological space, can be unified within the scope of the first resource diffusion from the environment to the organism, and P as the biological embedding starting point and the ending point of the first resource diffusion. This definition can be more clearly understood from the comparison between animals and fungi.

### 5 Mapping relationship between geometric conditions of n-simplex (n+1 vertices) and bio-geometric principles of biological space

In terms of methodology, this article only deals with the problem to match the number of simplex vertices and dimensionality. According to the principle of simplex topology geometry, it is only necessary to satisfy that the ℝⁿ space is composed of n+1 affine independent vertices (origin and standard basis), ensuring that the rank of the difference vector matrix is n (automatically satisfied by the origin method).

The general mapping relationship between this simplex construction condition and the biological geometry principles of biological space can be summarized as follows,

Firstly, the application of the origin method. Vertex d and vertex p serve as the geometric origins in the construction of D and D+1 simplices, respectively. Both parameters have deterministic topological geometric behavior origin connotations, which respectively define the starting embedding direction and starting position information in spatial embedding. The starting point condition for spatial embedding of plant populations does not satisfy the origin method, but under the condition that the height *h* of characteristic individuals remains constant, the origin method condition can still be satisfied by coordinate system translation.

Secondly, n+1 vertices are affine independent. Affine independence is specifically constrained and defined by the biological geometric sources of vertices, and the unified use of three-dimensional spatial coordinate reference systems and scale constraints for the macroscopic topological structure analysis of metabolic networks in this article, are prerequisites for ensuring vertex affine independence in the construction of biological spatial simplices.

## III Simplex model of the six modes of EMS

### 1 Individual-internal metabolic model (animal)

**Fig. 5.**
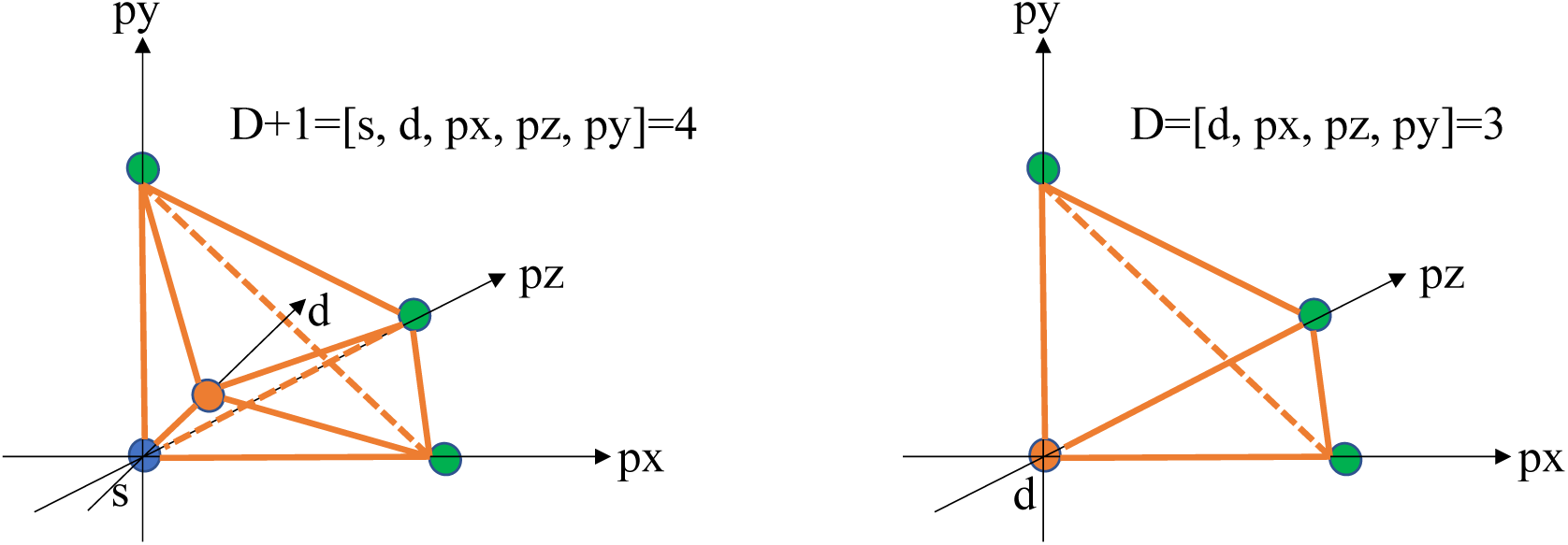
spatial embedding of individual-internal metabolic model (animal): Construction of biological spatial simplex. (1) Composition of vertices: **s**, resource space points (out of body; topological transformation of **p**); **d**, through the connection with s, the behavior space of resource capture (occupation) under phagocytosis is defined, which can be specifically defined as resource space point (in body); **px**, *x* coordinate points of three-dimensional space in body; **py**, *y* coordinate points in body; **pz**, *z* coordinate points in body. (2) Definition of simplex: D+1 simplex is defined by five affine independent vertices [s, d, px, py, pz] as a 4-simplex (with s as the origin). D-simplex is defined by four non coplanar vertices of [d, px, py, pz] (d is the origin). It is suitable for animals with internal circulation system development with phagocytic nutritional behavior.

The spatial embedding of animal individuals is a “resource point s” (embedding starting point) - resource capture (occupation) of resource point from outside to inside (d) - metabolic network construction of resource diffusion (utilization) in three-dimensional space (px, py, pz) in anima body. The d-s connection represents the resource possession (capture) activity of individual animals in the three-dimensional resource space (a one-dimensional resource diffusion activity). The engulfed resource points in the organism then spread to the whole body through the internal circulation system, which is equivalent to the three-dimensional reduction of the resource point space and full dispersion in the three-dimensional organism (a three-dimensional resource diffusion activity).

Taking the three-dimensional resource space as the reference frame, the construction of biological space is topologically represented as a four-dimensional “space expansion” mode defined by resource diffusion activities. Therefore, the so-called four-dimensional organism hypothesis adds a dimension of resource diffusion activity defined by the behavior of resource capture (phagocytosis) to the three-dimensional resource diffusion activity in vivo under the topological and geometric framework of ecosystem metabolic network.

In the simplex construction mode of origin method, [s, d, px, py, pz] five vertices have clear biological geometric meaning and topological spatial position relationship (s is in vitro; d is the center point of three-dimensional space in the body; px, py, pz are the three coordinate vertices defining the body space), with s as the origin, constraining the remaining four vertices to be in four linearly independent vector spaces respectively, and strictly satisfying the rank=n of the difference vector matrix composed of these n+1 parameters. [d, px, py, pz] also satisfies the above conditions.

### 2 Population-modular self-thinning model (plant)

**Fig. 6.**
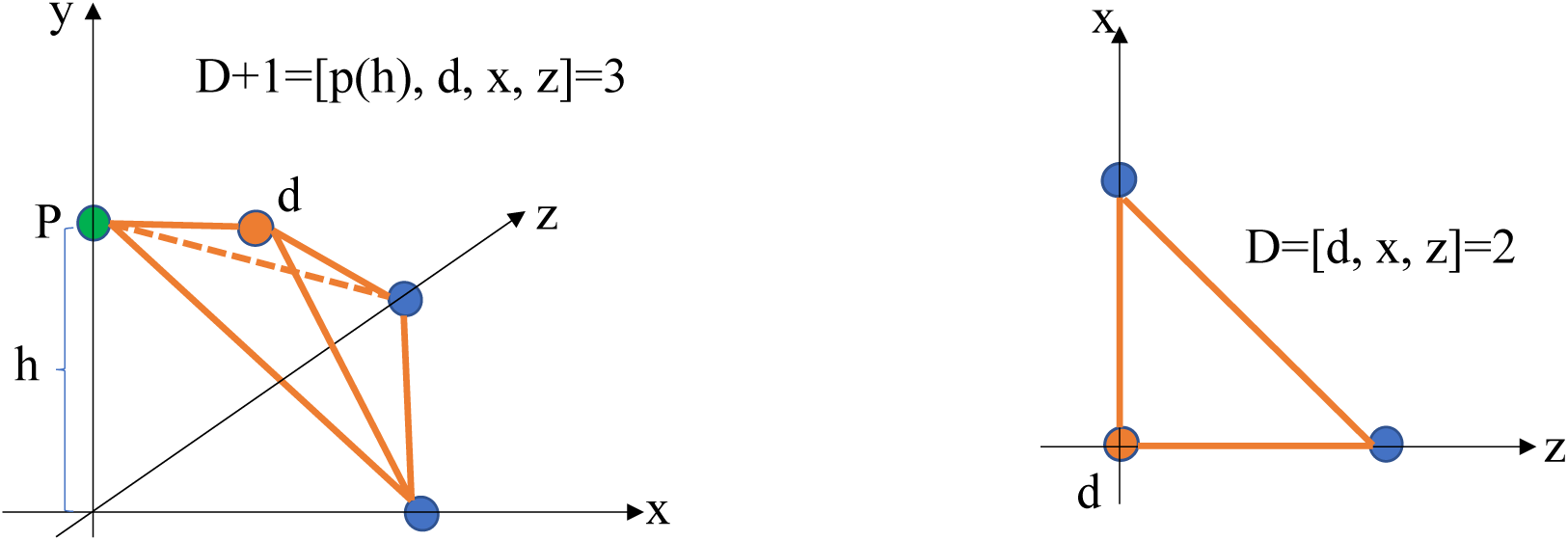
spatial embedding of population-modular self-thinning model (plant): Construction of biological spatial simplex. (1) The composition of the vertex: **p(h)**, the embedding starting point, indicating the initial position of the fixed growth individual in the three-dimensional resource space (the position on the y axis is defined by the constant value of the individual height, h); **d**, through the connection with p (h), the direction of spatial embedding (growth) in the process of self-thinning is defined, that is, the behavior space of resource capture (occupation); **x**, x coordinate vertex of 3D resource space; **z**, z coordinate vertex of 3D resource space. (2) Definition of simplex: D+1 simplex is defined as a 3-simplex (with p (h) as the origin) by four non coplanar vertices of [p (h), d, x, z]. D-simplex is defined by three non collinear vertices of [d, x, z] (d is the origin). multicellular plant with fixed growth suitable for the mode.

The spatial embedding of characteristic individuals in plant populations is a metabolic network construction constrained by self-thinning trajectories, which is “the embedding starting point p (p) defined by the constant individual height h -- the resource capture (occupation) defined by the d-p (h) perpendicular to the y axis -- and the two-dimensional resource diffusion (utilization) defined by the x and z axes”. That is, the characteristic individuals are embedded in the three-dimensional space in a fixed growth state, and the resources are captured (canopy extension) along the direction perpendicular to the y axis while maintaining the individual height h, supporting the resource diffusion (utilization) activities of a two-dimensional resource space (x, z). Among them, due to the neighborhood effect between individuals under crowded growth conditions, the reference function of the coordinate system of the y axis for individual resource utilization is lost. This is consistent with the bio-geometric principle of self-thinning under the conservation of metabolism. Therefore, taking the three-dimensional resource space as the reference frame, the dimension of effective metabolic activity is compressed to two dimensions (i.e. resource diffusion dimension D). As a fixed growth organism, the distribution of characteristic individual mass, that is, the construction of biological space, is still a three-dimensional process (i.e., the dimension of metabolic network D+1). This is a “piecemeal” construction mode of characteristic individual biological space.

Based on **h** as a constant greater than zero, the origin method condition of simplex geometry derivation (p as the origin) can be satisfied by coordinate translation. In this process, constrained by the above bio-geometric principles, h as a constant greater than zero and the condition that d-p transverse is greater than zero (and parallel to the x-z plane), the geometric conditions of simplex construction of plant population characteristic individuals [p(h), d, x, z] with four points not coplanar, and [d, x, z] with three points not collinear are jointly ensured.

### 3 Community-niche model (unicellular eukaryote)

**Fig. 7.**
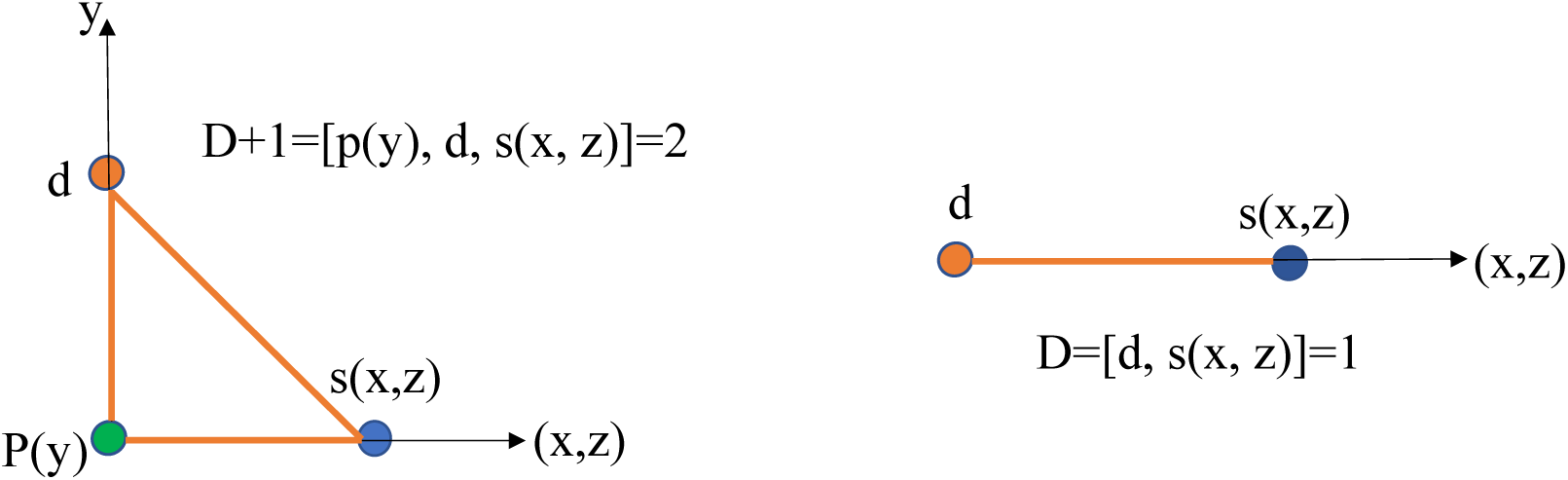
spatial embedding of community-niche model (unicellular eukaryote): Construction of biological spatial simplex. (1) The composition of the vertex: **p(y)**, the starting point of embedding (y indicates that the position of each species individual in the axis (niche axis) is fixed); **d**, the spatial embedding direction (parallel to y axis) is defined through the connection with P (y), that is, the behavior space of resource capture (occupation) specified by niche differentiation among species; **s(x, z)**, the individual activity space of the same species, is the resource diffusion (utilization) behavior space. (2) Definition of simplex: D+1 simplex is defined as a 2-simplex (with p(y) as the origin) by three non collinear vertices of [p(y), d, s(x, z)]. D-simplex is defined as 1-simplex (with d as the origin) by two non-coincident vertices of [d, s (x, z)]. unicellular eukaryote (diatom) suitable for the mode.

The unicellular eukaryote individual is topologically transformed into a point in the relative three-dimensional resource space, and its basic embedding (moving) behavior follows the diffusion activity mode. The construction of metabolic network is composed of “the embedding starting point P (y) with clear potential niche location -- niche differentiation and occupation of individuals of different species along the y axis (i.e. resource capture activities) -- secondary aggregation of individuals of the same species along the (x, z) axis (i.e. resource diffusion (utilization) activities)”. The embedding process can be abstracted as a trajectory formed by the point p(y) sweeping across the three-dimensional resource space along the p-d axis and the (x, z) axis, which is a two-dimensional convergent biological space construction process (a slice construction mode relative to the three-dimensional resource space). Compared with prokaryotes, single-cell eukaryotes in aquatic environments with larger cell size and more specific resource requirements, are more suitable for niche mode.

Under the condition of metabolic conservation, the individual activities in the default community are constrained by the full differentiation of species’ niche (the resource space is fully divided along the y-axis) and converge to the near two-dimensional resource space (defined by the x-axis and z-axis) corresponding to same species. Compared with plants, in the same two-dimensional activity space, the difference in topological characteristics of resource diffusion (utilization) activity space is determined by individual biological characteristics: subject to the non-fixed random diffusion activity mode, single-cell eukaryotes can be determined to appear in the coordinate plane, but their specific position in the plane is uncertain. Therefore, in topology, the whole is defined as one vertex s (x, z) or one coordinate axis (x, z).

Similarly, constrained by the above bio-geometric principles, the geometric conditions for simplex construction of characteristic individuals [p (y), d, s (x, z)] with three points not collinear and two points not coincident in three-dimensional resource space under the condition of metabolic conservation can be strictly guaranteed.

### 4 Community-neutral model (prokaryote)

**Fig. 8.**
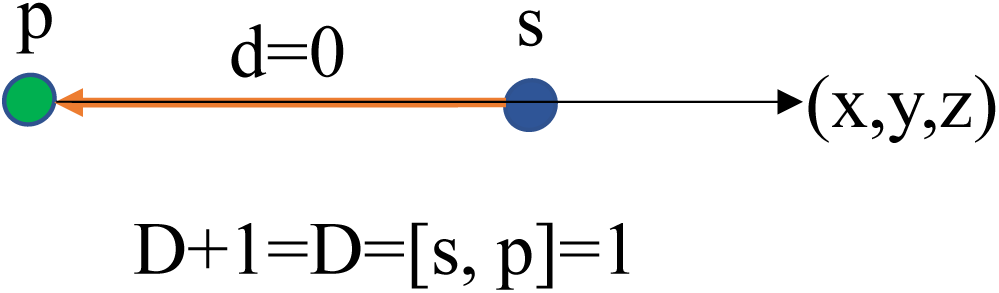
spatial embedding of community-neutral model (prokaryote): Construction of biological spatial simplex. (1) Composition of vertices: **p**, embedding unit; **s**, Embedded space. (2) Definition of simplex: two non-coincident vertices of resource space (s) and biological individual (P) form a 1-simplex, which jointly defines D+1 and D. prokaryote (bacillus) suitable for the mode.

d=0 means that the embedding activity of prokaryotes in the whole three-dimensional resource space conforms to the random diffusion mode (no definite direction), so the position reference value of the three-dimensional coordinate system of resource space in topology is lost, and the resource space is topologically transformed into a vertex (**s**)or a coordinate axis (x, y, z). Therefore, the parameter space defining the relative position relationship between P and S is only composed of [s, p], and the resource capture (possession) activity and resource diffusion (utilization) activity space are jointly defined by the one-way passive resource diffusion process driven by the environment. This is a 1-simplex defined by the linear topological connection between p and s, showing a linear biological space construction mode.

Compared with unicellular eukaryotes, bacteria in the water environment at the early stage of life evolution is more in line with this model (small size, high mobility, and more extensive resource demand (metabolism)), representing a highly simplified ecosystem organization model. Bacteria is an independent individual isolated from the environment, which strictly meets the geometric conditions of 1-simplex construction with two points [s, p] not coincident. In theory, the organizational nature of the population or community will not affect the above topological geometry principles.

### 5 Superbody system (fungal mycelium) and primitive soup system

**Fig. 9.**
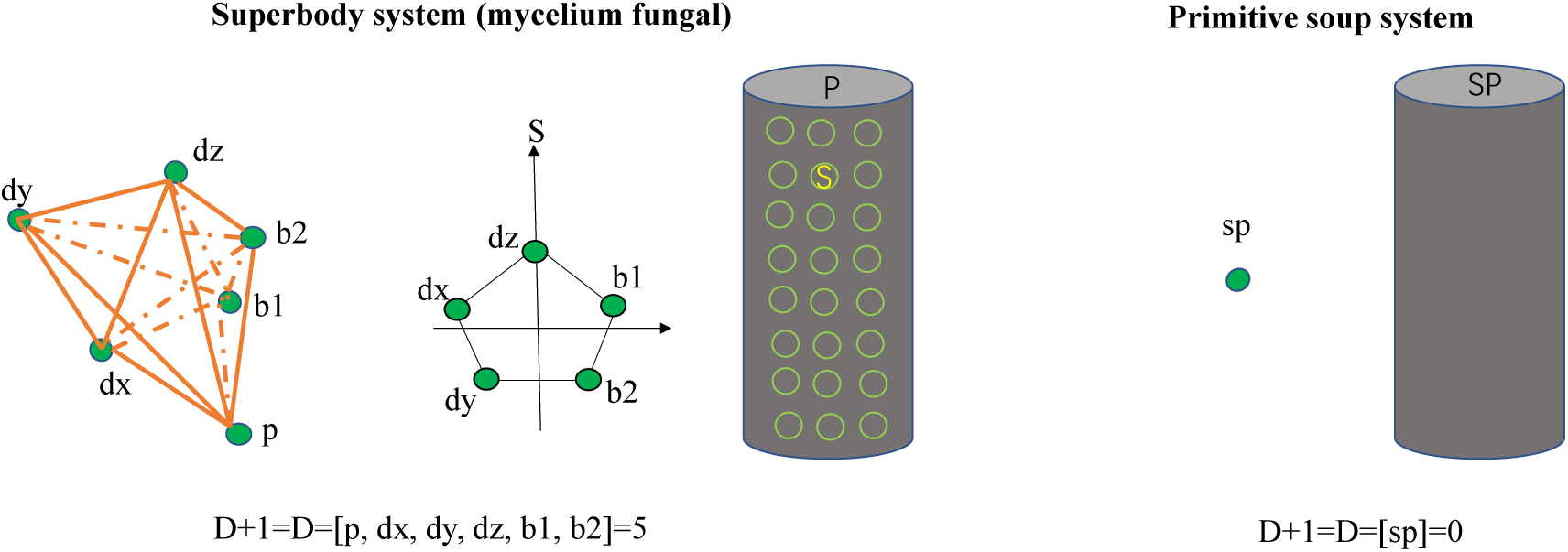
spatial embedding of superbody system (mycelium fungal) and primitive soup system: Construction of biological spatial simplex.

#### Biological space construction of superbody system (*f*(5)=5)

Mycelium fungi follow the topological construction of biological space named as “growth is metabolism”, by the form of three-dimensional network growth of mycelium. That is to say, the mycelium fully cuts and surrounds the resource space from the outside to the inside.

The biological space is constructed from five dimensions, including three physical space dimensions (determined by the network topological growth configuration, the resource capture (occupation) activity is defined by the three coordinate parameter dx, dy, dz), two biological diffusion dimensions (constrained by the surface area diffusion principle, and the resource diffusion (utilization) activity is defined by the two relative position parameters b1, b2 from the mycelium surface).

By transforming the minimum embedding space containing P and S into a two-dimensional projection plane (P-S), the topological configuration unit of mycelium metabolic network can be regarded as a pentagon structure, which directly reflects the resource capture and diffusion activities in five dimensions.

The biological space construction of mycelium fungi is similar to that of animals in terms of geometric principle, except that one-dimensional capture (phagocytosis) activity is replaced by three-dimensional networking growth, which represents the full occupation of three-dimensional resource space by mycelium fungi in a strict sense.

Since the relative internal and external position relationship between mycelium fungi and resource space is reversed thoroughly, the parameter space describing the relative position relationship between P and S needs to include the origin p, so D=D+1=5. Constrained by the biological geometry principle of biological space construction, the geometric condition of constructing 5-simplex with six vertices affine independence is strictly satisfied.

#### Biological space construction of primitive soup system (*f*(0)=0)

In primitive soup system, life components and environment are inseparable in structure.

Topologically, P and s are regarded as a 0-simplex (SP) composed of one point, so D+1=D=0.

## IV Transformation from topological geometry model to metabolic scaling model

First of all, through the unification of topological geometry model and metabolic scaling model in the internal metabolic model (animal; D=3, D+1=4, B∝M*^b^*=M^3/4^=M^D/D+1^), according to the metabolic steady-state equivalent transfer principle generally constructed in this paper on the rules of ecosystem organization, it is assumed that all organizational models follow the same conversion relationship between topological geometry principle and metabolic proportion biogeometry principle, that is, *b*=D/D+1, and the metabolic scaling model can be directly constructed,

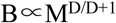

Secondly, under the constraint of the conservation of population metabolism (NB∝M^0^), taking the three-dimensional coordinate dimension of resource space as the scale constraint and the reference of relationship transformation, the following geometric transformation operation relationship can be gradually derived,

### 1 Metabolic network dimension D+1 and characteristic individual mass dimension D_M_

According to the definition of biological space, space embedding trajectory is topologically equivalent to the mass distribution process of characteristic individuals following specific organization rules and geometric principles in three-dimensional resource space, and its mass dimension D_M_ is equivalent to metabolic network dimension D+1. From this, it can be deduced,

Modular similarity system D_M_=D+1=3,

Niche system D_M_=D+1=2,

Neutral system D_M_=D+1=1.

### 2 Group mass distribution dimension D_NM_

The dimension (D_NM_) of group mass (NM) is defined by the macroscopic topological characteristics of the global embedding of the group as a unit in the three-dimensional resource space (Fig. 1).

In the three-dimensional resource space, the group embedding unit and the resource space are equivalent in topology. The global embedding trajectory is determined by the relative position relationship between them, which is only defined by one-dimensional distance parameters (such as internal and external relations), therefore D_NM_ =1, suitable for modular similarity system and niche system.

For the neutral model, the group is composed of a random set of individuals whose interactions between individuals is ignored. The relative position relationship between the group and resource space is uncertain, so D_NM_ = 0.

### 3 The functional relationship NM∝M^a^

Based on D_NM_ and D_M_, it can be deduced,

Modular similarity system NM∝M^1/3^,

Niche system NM∝M^1/2^,

Neutral system NM∝M^0^°

### 4 The functional relationship B∝M^b^

Based on NB∝M^0^, NM∝M^a^, it can be deduced,

Modular similarity system B∝M^2/3^=M^D/D+1^,

Niche system B∝M^1/2^=M^D/D+1^,

Neutral system B∝M = M^D/D+1^。

