## SUPPLEMENTAL TABLE 1 for "*EMS* hypothesis of Macroevolution"

### Supplementary materials

**Table S1 Macroevolution relationships of first resource diffusion, biological nodes, and second resource diffusion in ecosystem metabolism network**

| **Organizational model** | **Rules of ecosystem organization** | **natural selection (targeted at ecosystem)** ⇨ | **First resource diffusion (external metabolic network)** ⇨ | **Biological node p (status and configuration)** | ⇦ **Secondary resource diffusion (internal metabolic network)** | ⇦ **Complexity self-organization (taking life as the object)** | **Correlation in macroevolution** |
|---|---|---|---|---|---|---|---|
| Molecular organization model | Primitive soup system | Thioester world hypothesis: the molecular mechanism of metabolic network breeding is determined by the form of energy enrichment. | Functional network without macro structure (0,0) | Primitive soup system (RNA world hypothesis) | Chemical reaction network. | Breeding of metabolic function: chemical self-organization behavior dominates. | The unity of opposites between genetic priority and metabolic priority (Singh et al., 2025). |
| Group organization model | Neutral system (prokaryote) | Phylogenetic aggregation: the maximization of group living income under resource scarcity/harsh habitats drives small individuals' ecological behavior and phenotypic assimilation selection. | One dimensional network (1,1): one dimensional resource diffusion (environment) driven. | Ancestor (f): the kingdom that can be classified into three categories only by morphology. Bacilli are dominant. | Randomized network: highly flexible. Self organized evolution of metabolic modules at subnetwork scale. | Breeding of network structure: various metabolic modules start breeding in the process of random adaptation (plug and play, voyvodic et al., 2019) | Time is everything: mitochondrial precursors and chloroplast precursors need time to emerge in random, and then solidify after natural selection. |
| | Niche system (unicellular eukaryote) | Phylogenetic divergence: organisms as an environment, drive the ecological behavior and phenotypic differentiation of large individuals, and ensure the overall resource utilization efficiency. | Two dimensional network (1,2): driven by one-dimensional niche differentiation. | Ancestor (m): the most diverse and complex configuration differentiation. A plasticine experiment stage. **Diatoms** are the most representative. | Structured network: the foundation of network scale self-organization structure centered on mitochondria. | First endosymbiotic event (Martin et al., 2017). | The embryonic form of modern ecosystem and the breeding of life diversity. |

| | | | | | | | |
|---|---|---|---|---|---|---|---|
| | modular similarity system (plant) | Born to light: the growth and organization mode of geometric similarity of component organisms determined by the distribution mode of light resources. | Three dimensional network (2,3): driven by two-dimensional resource utilization activities. | **Producer:** modular assembly of photosynthetic canopy and branch network. Modular growth is the most dominant in photosynthetic plants. | Hierarchical network: the specialization of network self-organization structure marked by chloroplasts. | Second endosymbiotic events (Martin et al., 2017). | |
| | Internal metabolic system (animal) | Life lies in movement: phagocytosis fundamentally reversed the mode of decentralized food resource utilization: from passive to active. | Four dimensional network (3,4): driven by three-dimensional diffusion in the body. | **Consumer:** phenotypic trade-off optimization between exercise feeding and internal circulation -- generalization of slender body type. | Compartmentalization network): specialization of network self-organization structure marked by internal circulation system. | Internal circulation system. | Through the specialized evolution of biological metabolism function, the structure and function of modern ecosystem are established. |
| Individual organization model | Superbody system (mycelium fungal) | Growth is metabolism: determined by the nature and distribution of organic matter resources, the mycelium network as a whole can be regarded as the topological inversion of the internal circulation system of animals. | Five dimensional network (5,5): driven by five dimensional resource acquisition and utilization activities. | **Decomposer:** individual networking--pentahedral structure of mycelium network topological unit. | Centralized network: an efficient transmission network driven by the growth center at the top of the hypha, in conjunction with the external metabolic network. | Mycelium network. | |


3 Martin W, Tielens A, Mentel M, et al. The physiology of phagocytosis in the context of mitochondrial origin. Microbiology and Molecular Biology Reviews 2017, 81(3): e00008-17.

4 Singh J, Thoma B, Whitaker D, et al. Thioester-mediated RNA aminoacylation and peptidyl-RNA synthesis in water. Nature 644, 28, 933-944 (2025).

Voyvodic PL, Pandi A, Koch M, et al. Plug-and-play metabolic transducers expand the chemical detection space of cell-free biosensors. Nat Commun 2019, 10: 1697.

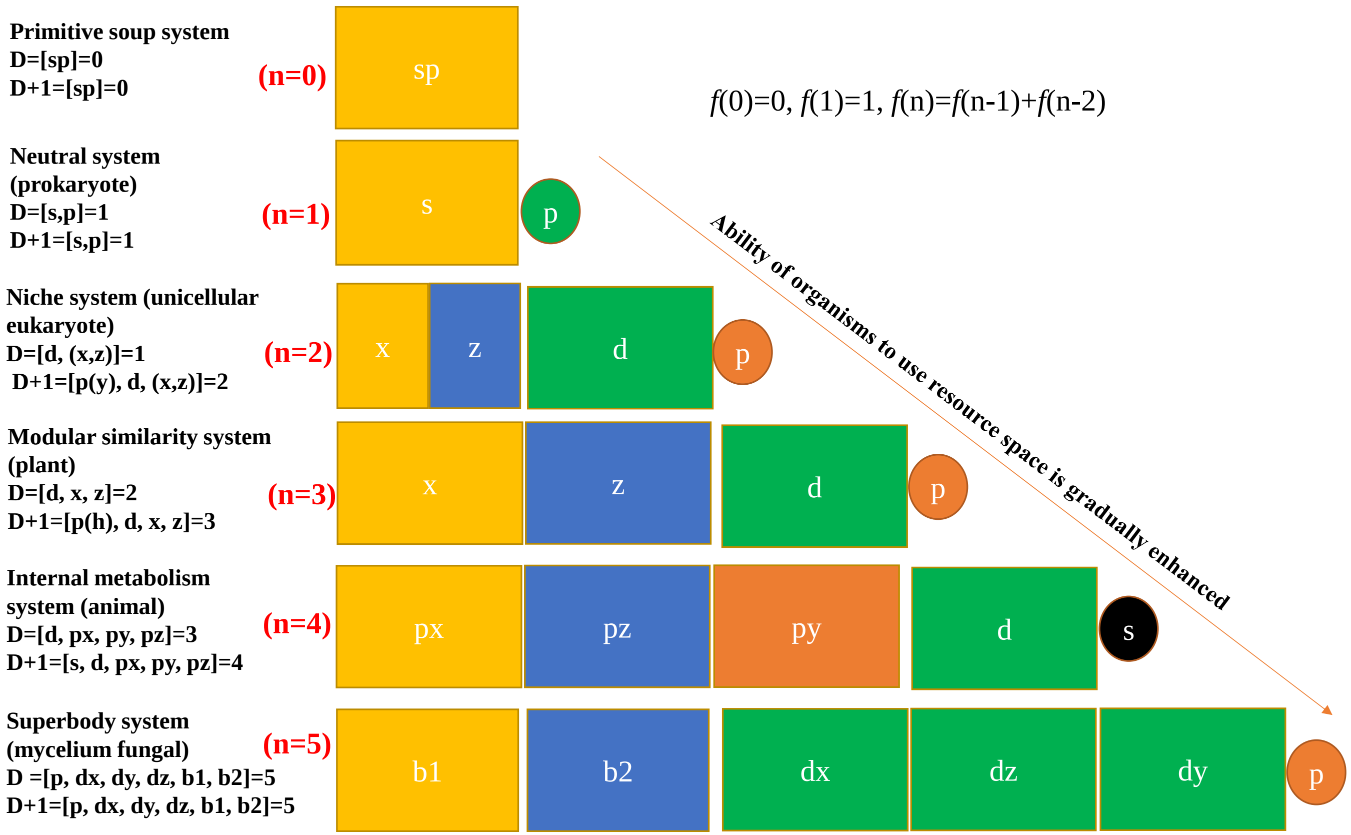


**Fig. S1 Origin and composition of the vertices of the D-simplex (resource diffusion dimension) and the D+1-simplex (metabolic network dimension).**
